# sFlt-1 (sVEGFR1) induces placental endoplasmic reticulum stress in trophoblast cell: implications for the complications in preeclampsia- an *in vitro* study

**DOI:** 10.1101/240481

**Authors:** Sankat Mochan, Manoj Kumar Dhingra, Betsy Varghese, Sunil Kumar Gupta, Shobhit saxena, Pallavi Arora, Neerja Rani, Arundhati Sharma, Kalpana Luthra, Sadanand Dwivedi, Neerja Bhatla, Rani Kumar, Renu Dhingra

## Abstract

**Background:** The concentration of sFlt-1, a major anti-angiogenic protein in maternal circulation has been seen to be raised in preeclamptic pregnancies. Endoplasmic reticulum (ER) stress represents one of the three (immunological, oxidative and ER stress) major stresses which placenta undergoes during pregnancies. The present study is designed to investigate the role of sFlt-1 in induction of ER stress in trophoblast cells.

**Materials and Methods:** Maternal serum levels of anti-angiogenic protein sFlt-1 and central regulator of unfolded protein response GRP78 was measured using sandwich ELISA. The expression of various ER stress markers (GRP78, eIF2α, XBP1, ATF6 and apoptotic protein CHOP) were analyzed depending on various treatments given to the trophoblast cells using Immunofluorescence, western blot and q-RT PCR.

**Results:** Increased expression of ER stress markers (GRP78, eIF2α, XBP1 ATF6 and apoptotic protein CHOP) was detected in the placental trophoblast cells treated with raised concentration of sFlt-1.

**Conclusion:** Significant upregulated expression of ER stress markers in trophoblast cells exposed with increased concentration of sFlt-1 suggested that it may be one of the anti-angiogenic factors present in maternal sera which not only contributes to oxidative stress but also may cause endoplasmic reticulum stress.

## Introduction

Hypertensive disorders of pregnancy are second to acute obstetric hemorrhage as a direct cause of maternal mortality and morbidity^1^ Preeclampsia (PE), the great obstetrical syndrome^2,3^ stands outs as a major subset of hypertensive disorders affecting every tenth^4^ pregnancy in the contemporary world in otherwise healthy pregnant women^1–3^. Placental malfunction has long been implicated in the pathogenesis of preeclampsia. A variety of proangiogenic (VEGF, PlGF) and antiangiogenic factors (sFlt-1, sEng) are elaborated by the developing placenta, and the balance among these factors is vital for normal placental development^5^. Increased production of anti-angiogenic factors disturbs this balance and results in the manifestation of the syndrome^5^. The entity of distinguished anti-angiogenic molecule sFlt1 (soluble -fms like tyrosin kinase-1) / (sVEGFR1) is a very conversant fact in preeclamptic pregnancies. sFlt-1, a spliced variant of receptor VEGFR1, is a potent circulating^6^ antagonist of VEGF and PlGF^7,8^. It is released from the placenta under a variety of “stress” conditions and is strongly implicated in the maternal syndrome of preeclampsia^9^. The increase in sFlt-1 concentration also occurs in normal pregnancies but, during the later parts of gestation whereas in preeclampsia this raised concentration is abnormal and happens too early in gestation, the underlying pathophysiology is still not discernible. The preeclamptic placenta experiences stress which has been variously categorized as immunological, oxidative and endoplasmic reticulum stress^9^. Amongst aforesaid, the underlay for endoplasmic reticulum (ER) and oxidative stress is ischaemia/hypoxia–reperfusion^10,11^. Endoplasmic reticulum (ER), in eukaryotic cells, is a specialized organelle that orchestrates the synthesis, folding and transport of one-third of the proteins^12^. Altered ER proteostasis leads to agglomeration of unfolded and/or mis-folded proteins in the ER lumen which activates the unfolded protein response (UPR)^13^ comprised of three highly conserved signaling pathways: The PERK–eIF2α pathway, which attenuates nonessential protein synthesis; ATF6 and IRE1–XBP1 pathways, increasing folding capacity by upregulation of the molecular chaperones GRP78 and GRP94 and phospholipid biosynthesis^14^. Increasing evidence has indicated altered morphology of ER including dilation of the ER cisternae within the syncytiotrophoblast in preeclamptic patients ^15^. Also the loss of ER homeostasis along with ER stress-induced trophoblastic apoptosis may play role in placental pathophysiology leading to preeclampsia^16,17,18,19^. The circulating levels of sVEGFR1/sFlt-1 in preeclamptic women may influence multiple endothelial endpoints, many of which have been previously shown to be dysregulated in preeclampsia but its effect on endoplasmic reticulum stress markers has not been explored so far. It is in this background we planned this study to find out whether the sFlt-1/sVEGFR1 can set to ER stress and how does it equipoise between homeostatic and apoptotic signaling during the ER stress *in vitro*.

## Materials and Methods

***In vitro* experiments** were carried out to analyze the effect of sFlt1–1 on induction of ER stress in trophoblast cells (BeWo cells). The human choriocarcinoma cell line (BeWo) was procured from American Type Culture Collection (ATCC) and maintained in F-12 HAM’S nutrient medium supplemented with 10% fetal bovine serum, 100 U/ml pencillin, 100μg/ml Streptomycin. Cells were passaged with 0.025% trypsin and 0.01% EDTA.

The study was divided into five experimental groups depending on various treatments given to BeWo cells [normotensive (NT) serum-Group1; normotensive (NT) serum + recombinant sFlt-1 (re-sFlt-1)-Group2; recombinant sFlt-1 (re-sFlt-1)-Group3; tunicamycin treatment-Group4 and no treatment-Group5].

For this, normotensive, non-proteinuric pregnant women (n=10) without any other medical complications were enrolled from the antenatal clinic and the inpatient ward of the Department of Obstetrics and Gynaecology, All India Institute of Medical Science, New Delhi, India. Protocol of the study was approved by the institute ethics committee and written informed consent was obtained from all the enrolled women. 5 ml of venous blood was collected, centrifuged at 1200 RPM for 4 minutes, serum was separated and stored in aliquots at -80ºC. The serum samples were stored for the ELISA and cell culture experiments.

The levels of sFlt-1 and GRP78 were estimated in the serum of normotensive, non-proteinuric (control) pregnant women by sandwich ELISA (sFlt-1 ELISA kit: R&D Systems Inc., Minneapolis, MN, U.S.A., GRP 78: Enzo Life Sciences, Inc.).

After the various treatments, activation of ER stress markers (GRP78, eIF2α, XBP1, ATF6 and CHOP) were assessed at various time points (8h, 14h, 24h) at protein (Immunofluorescence, Western blot) and gene level (qRT-PCR).

**Immunofluorescence Microscopy:** BeWo Cells were trypsinized, seeded, allowed to grow on coverslips in multiple well chamber and incubated at 37°C in 5% CO_2_. After 8h, 14 h and 24 h, cells were taken out from incubator and washed with PBS, fixed in 4% PFA for 15 min at room temperature. Cells were washed with PBS and permeabilized with PBS + 0.1% Triton X-100 followed by PBS washing. Nonspecific blocking was done using 5% normal goat serum in PBS and Triton X. Cells were incubated for 12 hours at 4°C in primary antibodies (anti GRP78 antibody (1:1000), anti eIF2α antibody (1:200), anti XBP1 antibody (1:200), anti ATF6 antibody (1:1000) and anti DDIT3/CHOP antibody (1:500). Cells were washed with PBSTx and thereafter incubated in secondary antibody in 1:500 dilution for 1 hour at room temperature in dark room. Cells were washed in PBS and mounted with flouroshield mounting media with DAPI on the slide and observed under the fluorescence microscope (Nikon Eclipse Ti-S elements using NiS-AR software).

**Western blot analysis:** Cells were lysed in SDS-PAGE sample buffer [2% SDS, 60 mM Tris-HCl (pH 6.8), 10% glycerol, 0.001% bromphenol blue, 0.33% mercaptoethanol] and boiled for 5 min. The lysates were analyzed by immunoblotting using 1:1000 of anti GRP78, 1:500 of anti eIF2α, anti XBP1, and 1:1000 of anti ATF6, anti CHOP (Abcam) for 12 h at 4°C. The blots were then incubated in secondary antibody (HRP conjugated) for 2 hours. The blots were visualized by treating the membranes in DAB, Tetrahydrochloride and H_2_O_2_. β-actin was used as protein loading control. The blots were then scanned in a gel documentation system, using Quantity 1 software (Bio-Rad, Hercules, CA, USA) for densitometric analysis.

**qRT-PCR (quantitative Real Time-Polymerase Chain Reaction):** RNA extraction was done from treated cells via Ambion (Invitrogen) kit followed by c-DNA conversion by Thermo revert aid H-minus reverse transcriptase kit. cDNA was amplified by quantitative RT-PCR for determining mRNA expression of GRP78, eIF2α, XBP1, ATF6 and CHOP against gene of interest with an internal control (β actin and GAPDH).

### Statistical Analysis

Data was analyzed by Microsoft Office Excel Version 2013 and Graph Pad Prism. For mRNA expression by qRT-PCR, relative quantification cycles of gene of interest (ΔCq) was calculated by ΔCq = Cq (target) - Cq (reference). Relative mRNA expression with respect to internal control gene was calculated by 2^-ΔCq^. Data was presented in mean±SD/median (Range) as appropriate. Average level of the variable between the two groups was compared by paired t-test/Wilcoxon signed rank test. For comparing more than two groups, ANOVA with Bonferroni’s multiple comparison test/Kruskal Wallis with Dunn’s multiple comparison test were used. *p* value<0.05 was considered statistically significant.

## Results

The mean systolic and diastolic blood pressures were 117.8 ± 7.34 mm Hg (mean ± SD) and 74.2 ± 6.39 mm Hg (mean ± SD) respectively. The body mass index was 23.67 ± 3.59 (mean ± SD). Urine protein was analyzed by urine dipstick method and was found either in trace or nil.

**Levels of sFlt-1 and GRP78 in normotensive non-proteinuric pregnant women:** The levels of sFlt-1 and GRP78 in the maternal serum were found to be 2187.67 + 92.64 pg/ml and 970.75 + 107.78 ng/ml respectively.

### *In Vitro* Experiments

**Weaker expression of GRP78, eIF2**α**, XBP1, ATF6 and CHOP was observed in normotensive sera (NT) treated BeWo cells [Figure 2a-c]:** Immunofluorescence microscopy revealed no significant expression of ER stress markers at different time points [Figure 2a] However time kinetics studies using immunoblot demonstrated higher expression of GRP78 and eIF2α at 14 h as compared to 8 h and 24 h whereas XBP1, ATF6 and CHOP expressions were found more at 24 h as compared to 8h and 14 h [Figure 2b]. mRNA levels of GRP78, eIF2α, XBP1, ATF6 and CHOP were found to be higher at 14 h as compared to 8 h and 24 h [Figure 2c].

**Figure 1:**
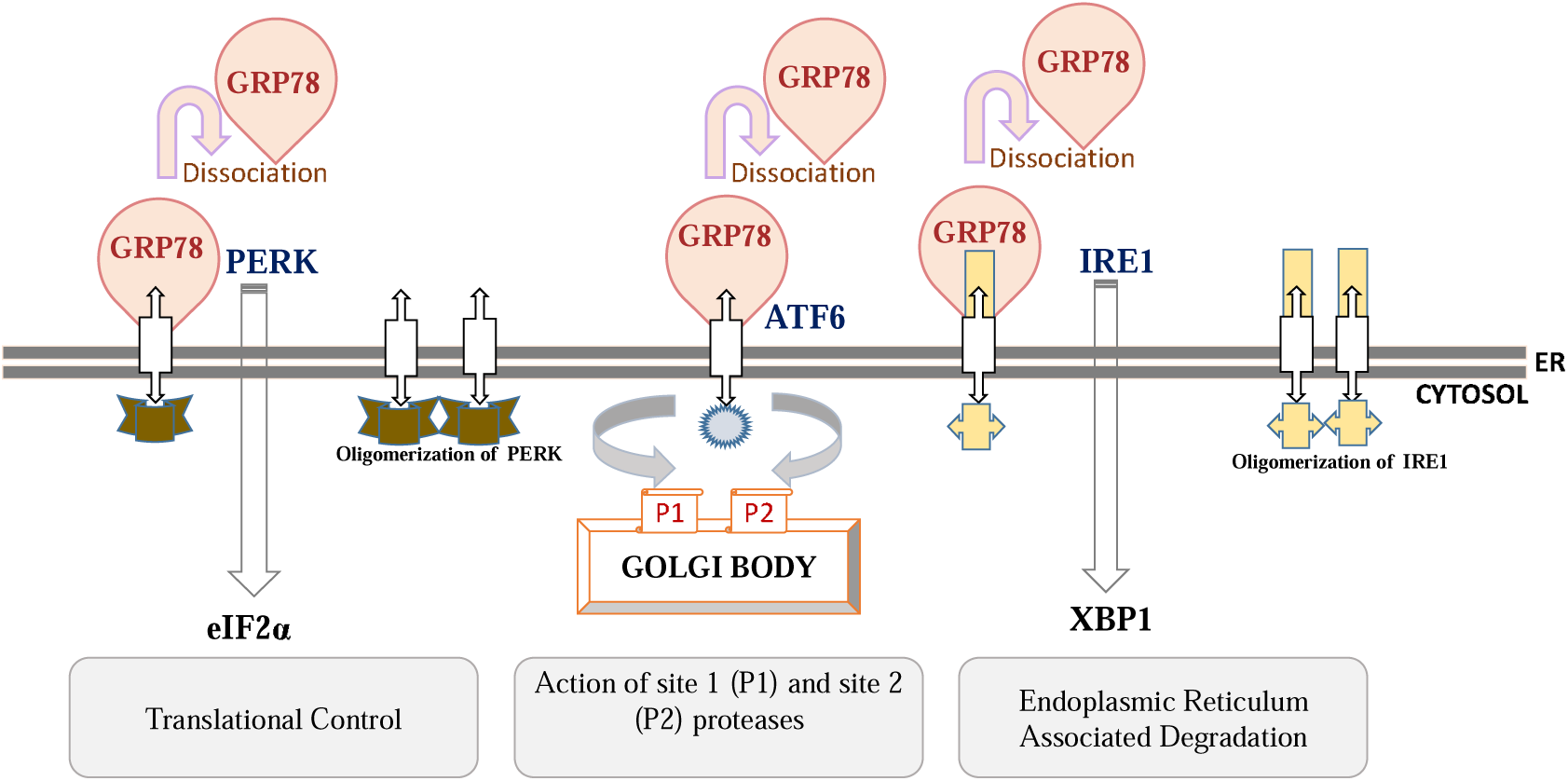
ER stress pathway

**Figure 2a:**
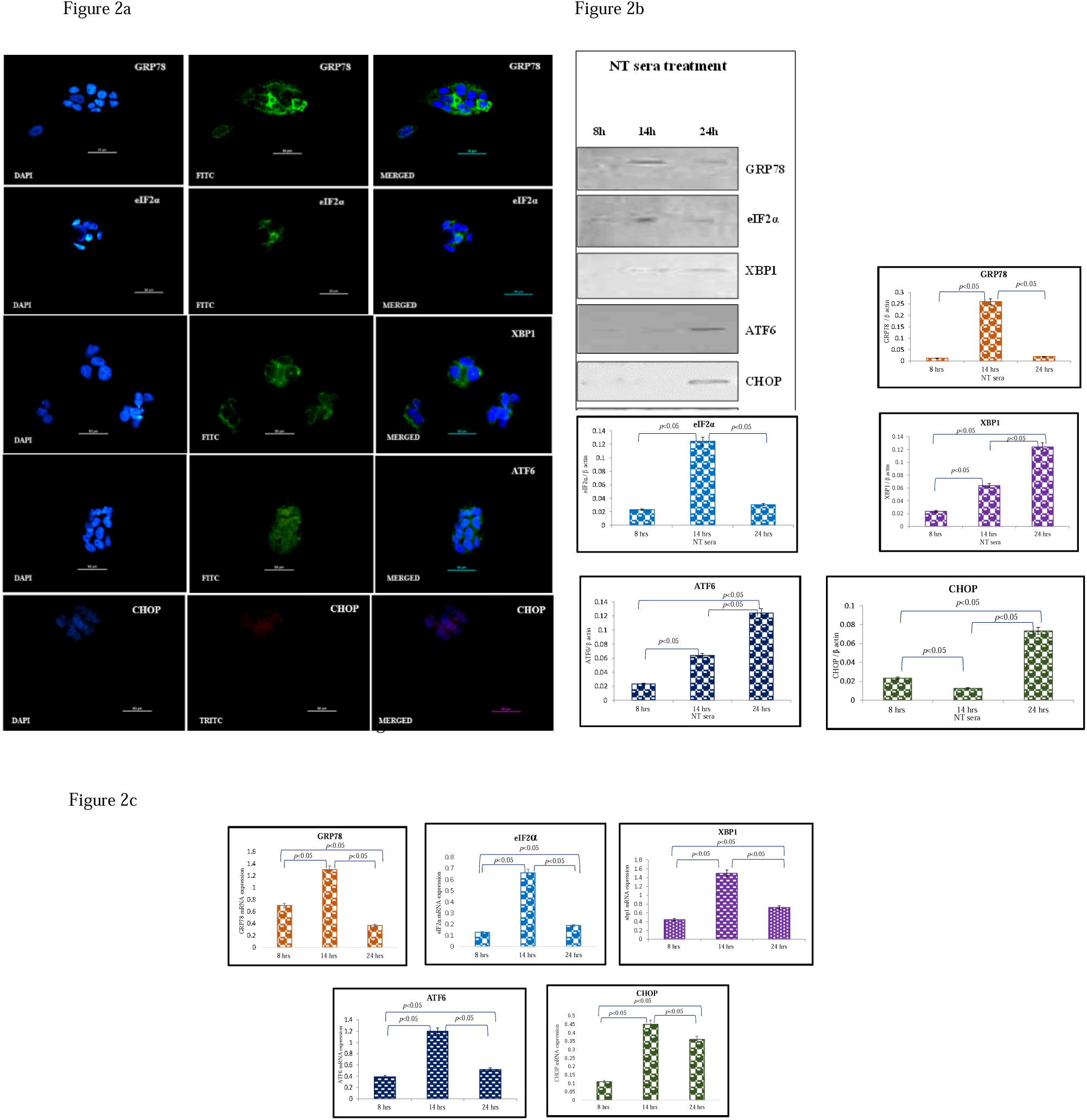
Representative immunofluorescence staining pattern of anti-GRP78 antibody positive BeWo cells following 8 h, anti-eIF2α, anti-XBP1, anti-ATF6 antibody positive BeWo cells following 14 h and anti-DDIT3 (CHOP) antibody positive BeWo cells following 24 h treatment of NT sera; Figure 2b: Representative images of immunoblot showing the expression of ER stress markers, GRP78, eIF2α, XBP1, ATF6 and CHOP in BeWo cells. β-Actin was used as protein loading control. The Bar diagrams represent the normalized values of the markers. Results are representative of 7 independent experiments. Data presented as mean ± SD. Statistical analysis was done using one way ANOVA with Bonferroni’s post hoc; Figure 2c: Bar diagrams represent the relative mRNA expression of GRP78, eIF2α, XBP1, ATF6 and CHOP and was found maximum at 14 h. GAPDH was used as positive control. Data presented as mean ± SD. One way ANOVA with Bonferroni’s post hoc test was applied (*p* values indicated on graph itself

**Significantly increased expression of GRP78, eIF2**α**, XBP1, ATF6 and CHOP in normotensive sera along with re-sFlt-1 treated BeWo cells [Figure 3a-c]:**

**Figure 3a:**
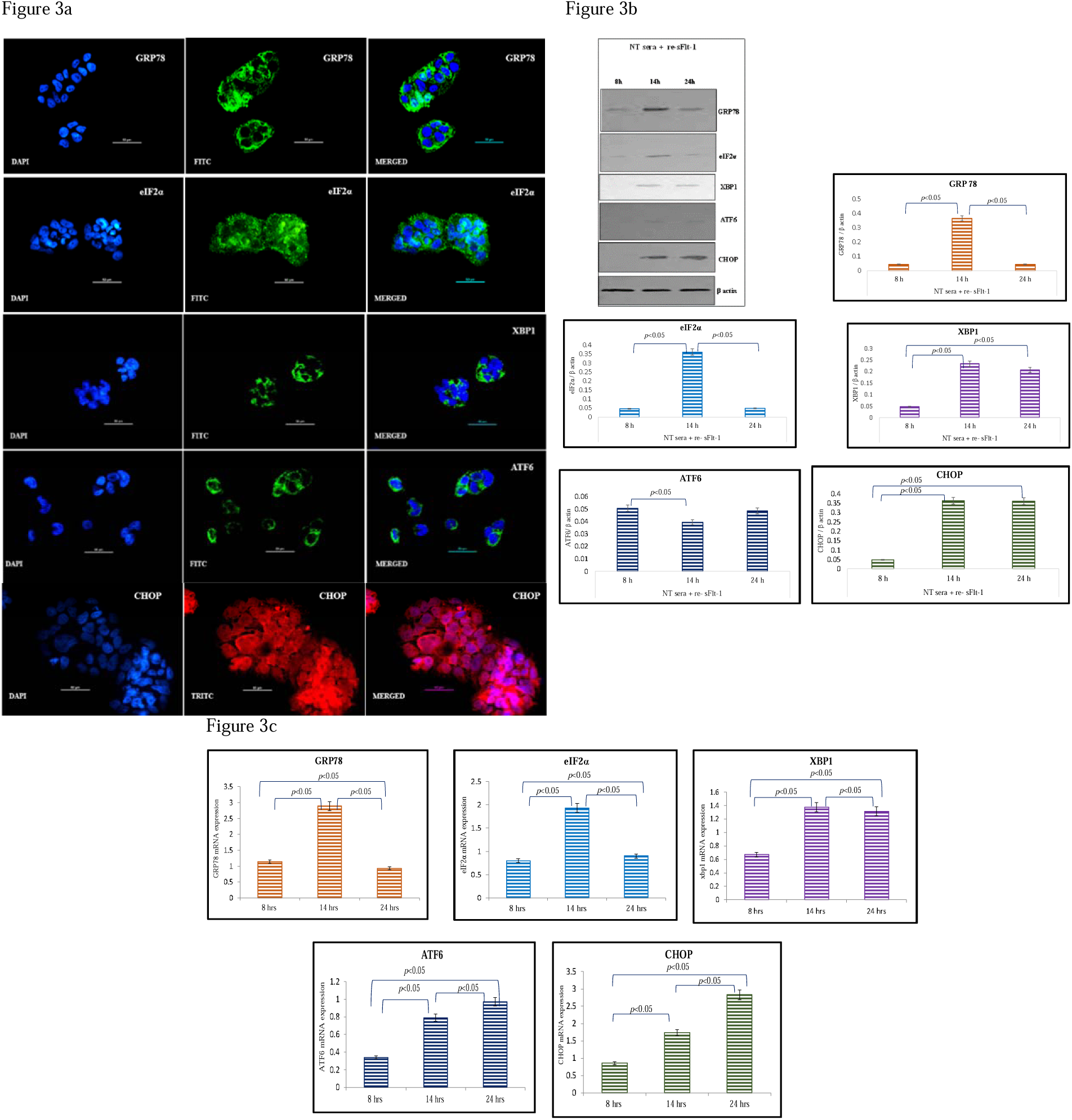
Representative immunofluorescence staining pattern of anti-GRP78 antibody positive BeWo cells following 8 h, anti-eIF2α, anti-XBP1, anti-ATF6 antibody positive BeWo cells following 14 h and anti-DDIT3 (CHOP) antibody positive BeWoo cells following 24 h treatment of NT sera + re-sFlt-1; Figure 3b: Representative images of immunoblot showing the expressionon of ER stress markers, GRP78, eIF2α, XBP1, ATF6 and CHOP in BeWo cells. β-Actin was used as protein loading control. Thehe Bar diagrams represent the normalized values of the markers. Results are representative of 7 independent experiments. Datata presented as mean ± SD. Statistical analysis was done using one way ANOVA with Bonferroni’s post hoc; Figure 3c: Barar diagrams represent the relative mRNA expression of GRP78, eIF2α, XBP1, ATF6 and CHOP. GAPDH was used as positiveve control. Data presented as mean ± SD. One way ANOVA with Bonferroni’s post hoc test was applied (*p* values indicated onon graph itself)

Immunofluorescence microscopy revealed significant expression of GRP 78 at 8 hours, eIF2α, XBP1, ATF6 at 14 hours and CHOP at 24 hours. Characteristic stress granules in case of eIF2α were observed at 14 hours [Figure 3a]. The expression of GRP78, eIF2α and XBP1 proteins were found to be higher at 14 h as compared to 8 h and 24 h time points. ATF6 expression was found to be higher at 8 and 24 h time points as compared to 14 h. Maximum expression of CHOP was found at 24 h [Figure 3b]. mRNA levels of GRP78, eIF2α and XBP1 were found to be higher at 14 h whereas ATF6 and CHOP mRNA levels were found to be higher at 24 h [Figure 3c].

**Expression of GRP78, eIF2**α**, XBP1, ATF6 and CHOP) was raised in re-sFlt-1 treated BeWo cells [Figure 4a-c]:** Significant expression of XBP1, ATF6 and CHOP was observed at 14 hours with 12 ng/ml concentration of re-sFlt-1 as compared to 9 ng/ml. Characteristic stress granules in case of eIF2α were also seen at 14 hours [Figure 4a]. Immunoblot revealed higher expressions of GRP78 and eIF2α at 14 h with 12 ng/ml concentration (c2) of recombinant sFlt-1 as compared to 9 ng/ml concentration (c1). Expression of XBP1 was found to be higher at 24 h with 12 ng/ml concentration (c2). ATF6 expression was found consistent at both time points (14 h and 24 h). CHOP expression was observed maximum at 24 h with 9 ng/ml (c1) concentration of recombinant sFlt-1 [Figure 4b]. GRP78 mRNA levels were found maximum at 14 h (c2) whereas mRNA levels of eIF2α, XBP1, ATF6 and CHOP were found maximum at 24 h (c2) [Figure 4c].

**Figure 4a:**
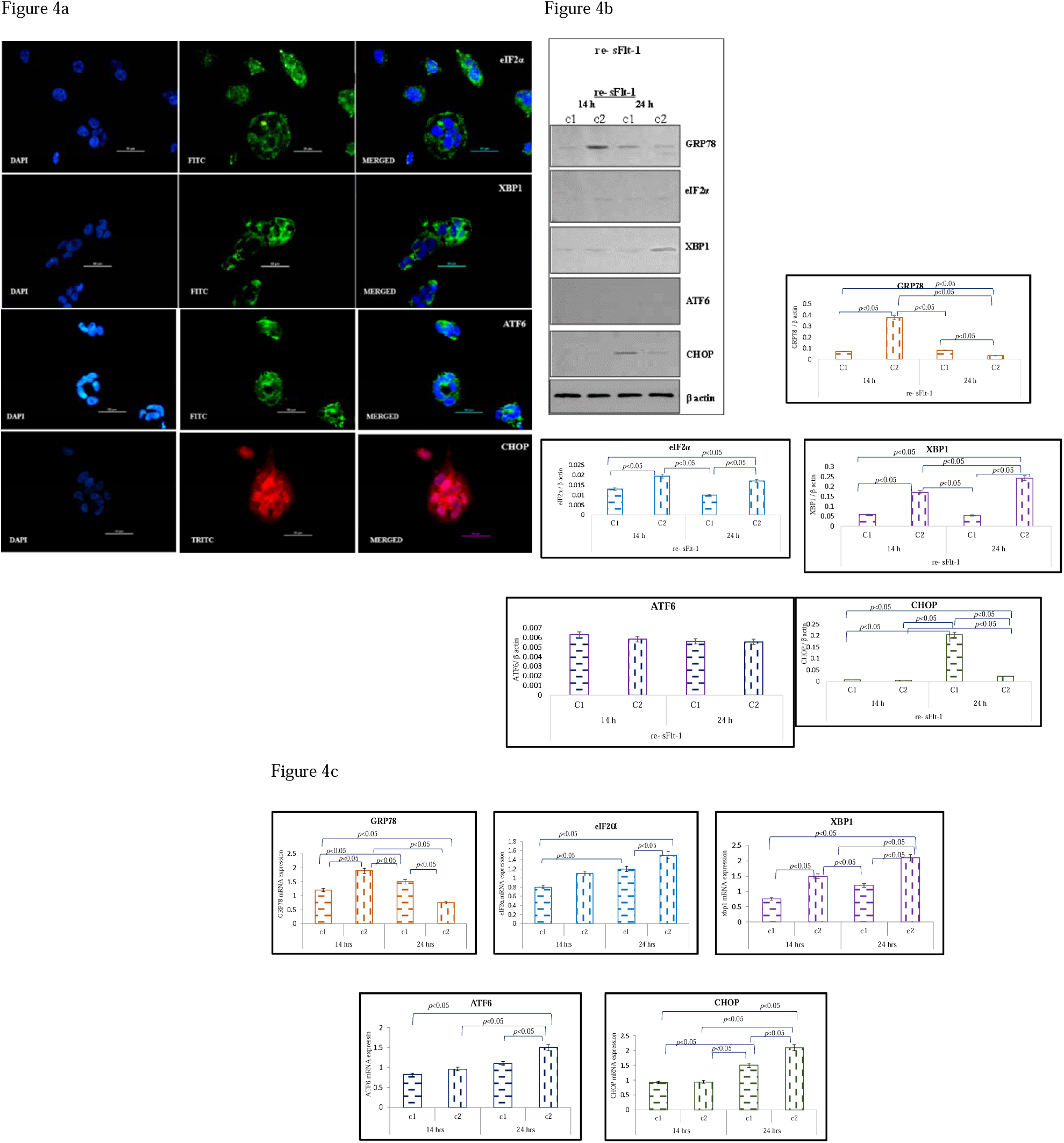
Representative immunofluorescence staining pattern of anti-eIF2α, anti-XBP1, anti-ATF6 and anti-DDIT3 (CHOP)P) antibody positive BeWo cells following 14 treatment of re-sFlt-1 (12ng/ml); Figure 4b: Representative images of immunoblott showing the expression of ER stress markers, GRP78, eIF2α, XBP1, ATF6 and CHOP in BeWo cells. β-Actin was used asasprotein loading control. The Bar diagrams represent the normalized values of the markers. Results are representative of 77 independent experiments. Data presented as mean ± SD. Statistical analysis was done using one way ANOVA with Bonferroni’s s post hoc; Figure 4c: Bar diagrams represent the relative mRNA expression of GRP78, eIF2α, XBP1, ATF6 and CHOP. GAPDHH was used as positive control. Data presented as mean ± SD. One way ANOVA with Bonferroni’s post hoc test was applied (*pp* values indicated on graph itself)

**Earmarked expression of GRP78, eIF2**α**, XBP1, ATF6 and CHOP in Tunicamycin treated BeWo cells [Figure 5a-c]:** The expression of all the markers were observed more so with 5 µg/ml dose of tunicamycin as compared to its lower dose [Figure 5a]. Immunoblot data revealed higher expressions of GRP 78 and eIF2α at 14h and 24 h as compared to 8 h. Expression was even more when its concentration was increased. XBP1, ATF6 and CHOP expressions was found to be higher with 5 µg/ml of tunicamycin as compared to 2.5 µg/ml of tunicamycin [Figure 5b]. GRP78, eIF2α, XBP1 and ATF6 mRNA levels were found to be higher at 14 h as compared to 8 h and 24 h. CHOP mRNA levels were found maximum at 24 h [Figure 5c].

**Figure 5a:**
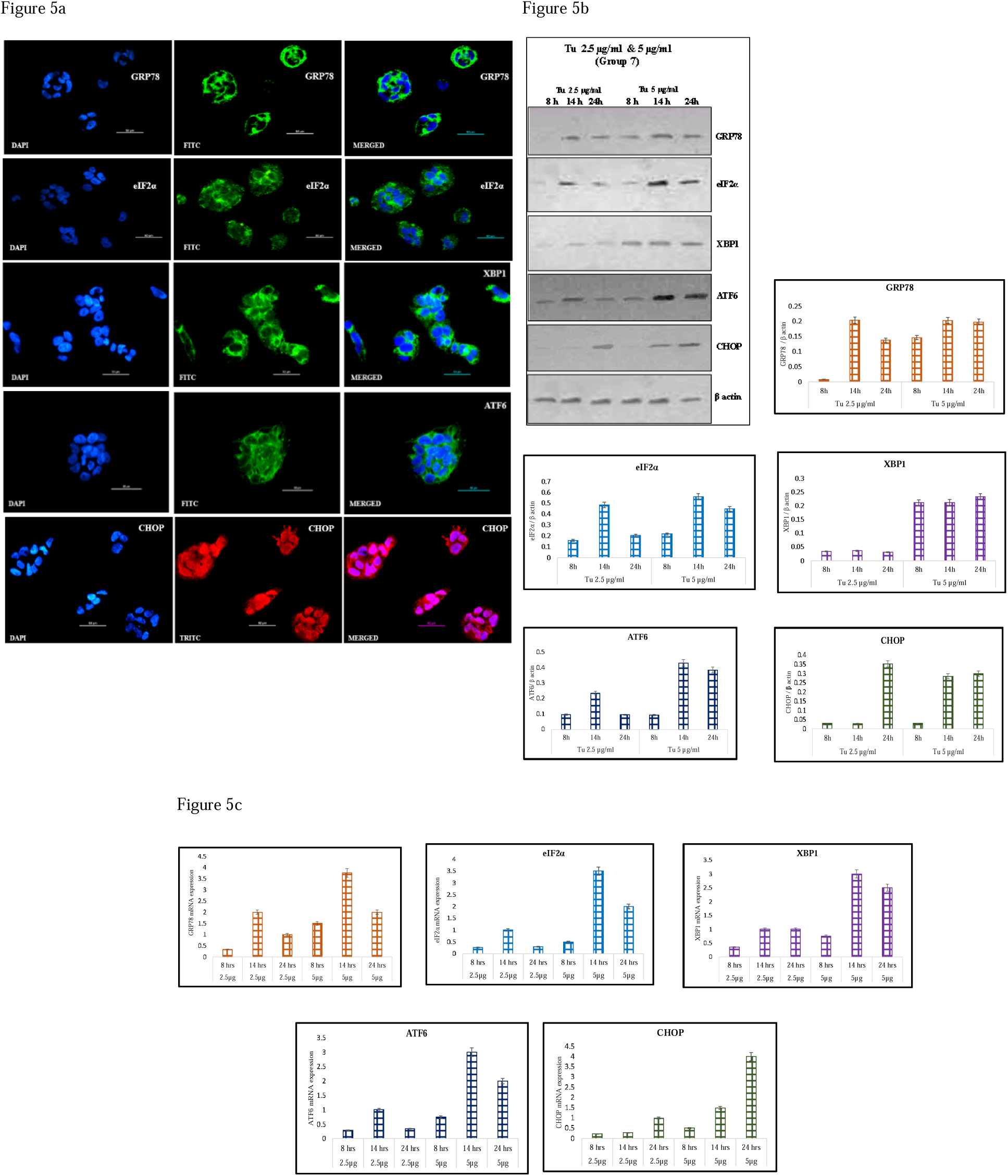
Representative immunofluorescence staining pattern of anti-GRP78 antibody positive BeWo cells following 8 h, anti-eIF2α, anti-XBP1, anti-ATF6 antibody positive BeWo cells following 14 h and anti-DDIT3 (CHOP) antibody positive BeWo cells following tunicamycin treatment (5µg/ml). Figure 5b: Representative images of immunoblot showing the expression of ER stress markers, GRP78, eIF2α, XBP1, ATF6 and CHOP in BeWo cells. β-Actin was used as protein loading control. The Bar diagrams represent the normalized values of the markers. Results are representative of 7 independent experiments. Data presented as mean ± SD. Statistical analysis was done using one way ANOVA with Bonferroni’s post hoc; Figure 5c: Bar diagrams represent the relative mRNA expression of GRP78, eIF2α, XBP1, ATF6 and CHOP. GAPDH was used as positive control. Data presented as mean ± SD. One way ANOVA with Bonferroni’s post hoc test was applied (*p* values indicated on graph itself)

**Feeble expression of GRP78, eIF2**α**, XBP1, ATF6 and CHOP in untreated BeWo cells [Figure 6a-c]:** Weak expression of GRP78 at 8 hours and eIF2α at 14 hours was observed. However, no expression of XBP1, ATF6 and CHOP was observed at different time points [Figure 6a]. Western blot analysis did show some expression of GRP78, eIF2α and XBP1 at 14 h as compared to 8 h and 24 h. ATF6 and CHOP expressions were found maximum at 24 h [Figure 6b]. GRP78, eIF2α and ATF6 mRNA levels were found maximum at 14 h however XBP1 and CHOP mRNA levels were found to be higher at 24 h [Figure 6c]. However the levels of these markers were lower as compared to tunicamycin, NT, NT+re-sFlt-1 and re-sFlt-1 alone treated BeWo cells.

**Figure 6a:**
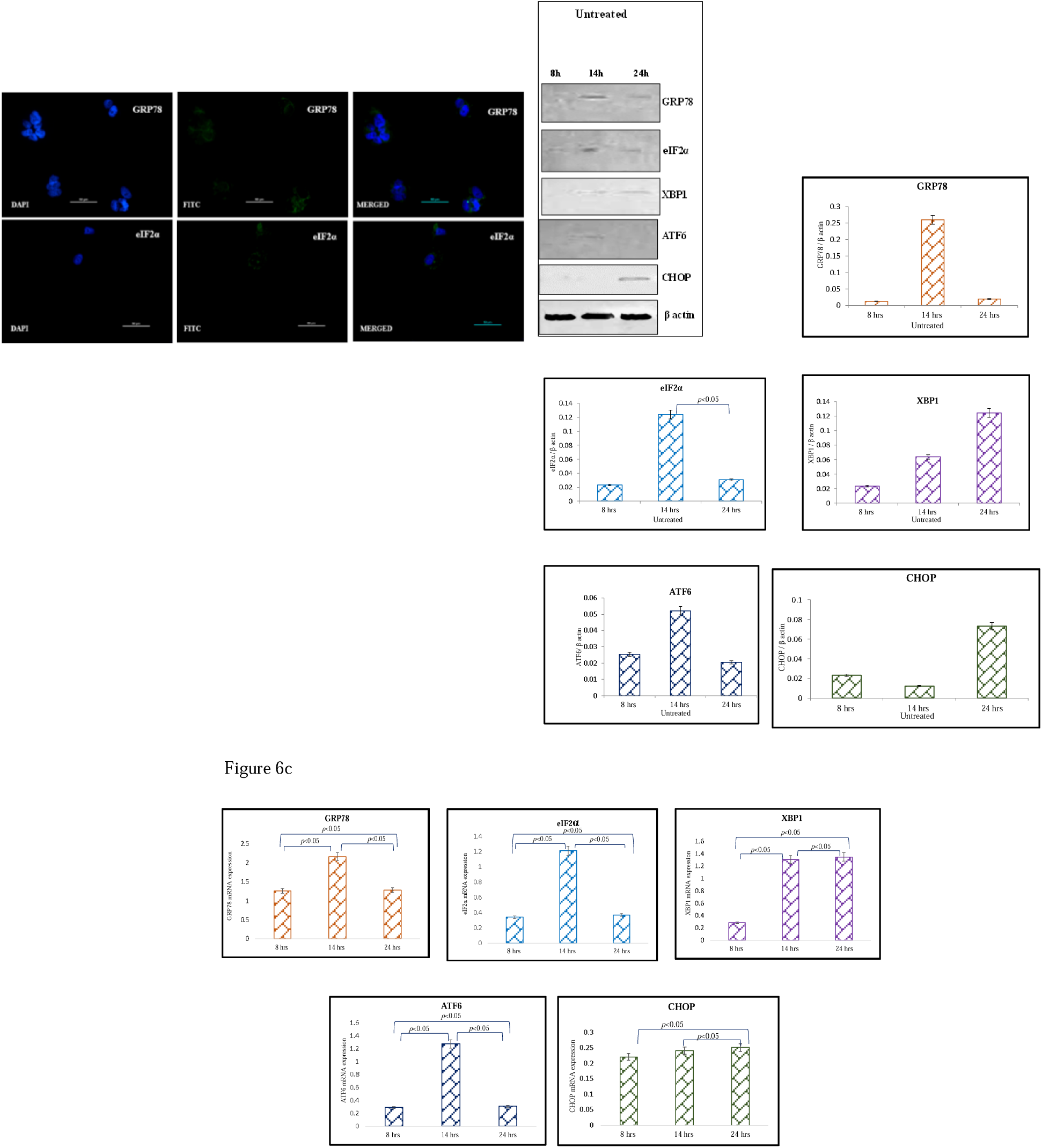
Representative immunofluorescence staining pattern of anti-GRP78 antibody positive BeWo cells following 8 h and anti-eIF2α antibody positive BeWo cells following no treatment. Figure 6b: Representative images of immunoblot showing the expression of ER stress markers, GRP78, eIF2α, XBP1, ATF6 and CHOP in BeWo cells. β-Actin was used as protein loading control. The Bar diagrams represent the normalized values of the markers. Results are representative of 7 independent experiments. Data presented as mean ± SD. Statistical analysis was done using one way ANOVA with Bonferroni’s post hoc; Figure 6c: Bar diagrams represent the relative mRNA expression of GRP78, eIF2α, XBP1, ATF6 and CHOP. GAPDH was used as positive control. Data presented as mean ± SD. One way ANOVA with Bonferroni’s post hoc test was applied (*p* values indicated on graph itself)

**Comparison of normalized protein values and mRNA levels of ER stress markers between NT sera and NT sera + re-sFlt-1 treated BeWo cells [Figure 7a-j]:** At 8 h, expressions of all ER stress markers were found to be higher in NT+ re-sFlt-1 treated BeWo cells as compared to NT sera and the difference was found to be statistically significant [GRP78 (*p*<0.0001), eIF2α (*p*<0.0001), XBP1 (*p*<0.0001), ATF6 (*p*<0.0001) and CHOP (*p*<0.0001)] [Figure 7a-e]. At 14 h, GRP78, eIF2α, XBP1 and CHOP expressions were also higher in NT+ re-sFlt-1 treated BeWo cells as compared to NT sera and the difference was statistically significant [GRP78 (*p*<0.0001), eIF2α (*p*<0.0001), XBP1 (*p*<0.0001) and CHOP (*p*<0.0001)] [Figure 7a-e]. At 24 h, expressions of all ER stress markers were found to be higher in NT+ re-sFlt-1 treated BeWo cells as compared to NT sera treated BeWo cells and the difference was statistically significant. GRP78 (*p*<0.0001), eIF2α (*p*=0.0002), XBP1 (*p*=0.0129), ATF6 (*p*=0.0001) and CHOP (*p*<0.0001) [Figure 7a-e]. At 8, 14 and 24 h, GRP 78, eIF2α, XBP1, ATF6 and CHOP mRNA levels were found to be higher in NT+ re-sFlt-1 treated BeWo cells as compared to NT sera treated cells and the difference was found to be statistically significant [GRP78 (*p*<0.0001), eIF2α (*p*<0.0001), XBP1 (*p*<0.0001), ATF6 (*p*<0.0001) and CHOP (*p*<0.0001)] [Figure 7f-j].

**Figure 7a-e:**
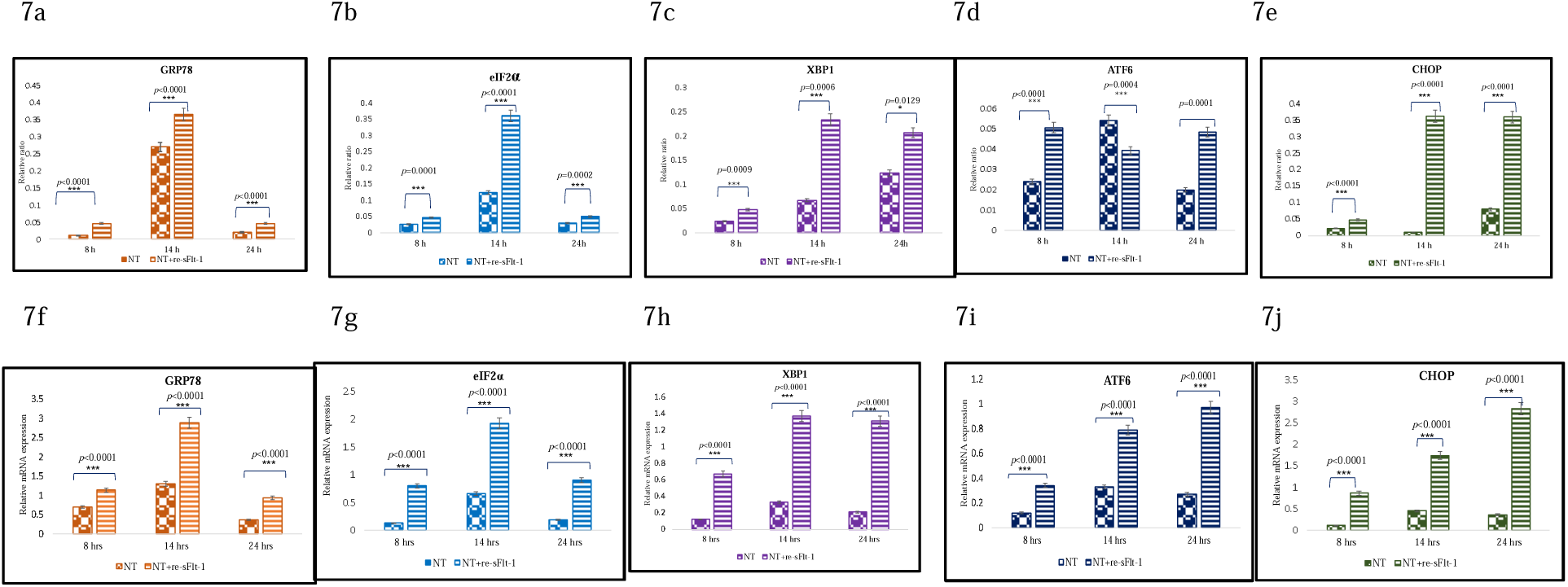
Comparison of normalized protein values of various ER stress markers between NT sera treated BeWo cells and cells which received both NT sera + re-sFlt-1; Figure 7f-j: Comparison of relative mRNA expression of ER stress markers between NT sera treated BeWo cells and cells which received both NT sera + re-sFlt-1. Statistical analysis was done using paired t test at all the indicated time points for all markers

## Discussion

The peculiarity of human placentation is egregiously concerned with the physiological remodelling of maternal spiral arteries through invading trophoblast to produce a low resistance and high capacitance uterine circulation^20^. Inaccuracy in this very event places the placenta at increased risk of ischemia–reperfusion type insult, a powerful stimulus for the generation of oxidative stress^21^ which leads to adverse obstetric outcomes; preeclampsia is one amongst them^22^. Defective placentation leads to release of a mixture of proteins that stimulate maternal endothelium and result in manifestation of the symptoms of disease. One such protein is the soluble form of the receptor, vascular endothelial growth factor receptor-1 (sVEGFR-1/sFlt1–1), which is produced in excess by the placenta of preeclamptic women in first trimester of ongoing pregnancy ^23^. The binding of sFlt1–1 to VEGF reduces concentrations of free VEGF in early gestation which is essential for effective vasculogenesis, angiogenesis, trophoblast cell proliferation and thereby affecting the normal placental development^24^. The resultant subsidence in bioavailability of VEGF makes placenta more prone to oxidative cell damage^25^. Thus, whenever serum levels of sFlt-1 increase, their binding with VEGF may reduce the circulating (free) VEGF levels below a critical threshold thereby adversely affecting the placental development and in turn, pregnancy^25,26,27^. In the patients of Indian origin, our group for the first time reported the alterations in the levels of anti-angiogenic (sFlt-1) and pro-angiogenic (VEGF, PlGF) factors in sera and PlGF in the urine of PE patients^28^. The sera of pregnant women (NT and PE) were later used for *in vitro* experiments. The BeWo cells subjected to treatment with preeclamptic sera showed increased amount of ER stress as compared to NT sera (unpublished results). There may be many factors differing in the two sera which could result in varying ER stress responses, so in the present study, we aimed to explore the role of sFlt-1 in inciting endoplasmic reticulum (ER) stress, in trophoblast cells (choriocarcinoma cells of placental origin-BeWo cells). We estimated the levels of sFlt-1 and GRP78 (central regulator of ER stress) in serum of 10 healthy pregnant women (normotensive, non proteinuric). The serum sFlt-1 levels in normotensive, non-proteinuric pregnant women were 2187.67 + 92.64 pg/ml. The BeWo cells (human choriocarcinoma cell line of placental origin which mimics *in vivo* syncytialisation of placental villous trophoblast) were subjected to various concentrations of sFlt-1 starting from lower concentrations as in normotensive sera, followed by normotensive sera along with re-sFlt-1 and administration of two dosages of recombinant sFlt-1; 9 and 12 ng/ml independently. The expression of ER stress markers (GRP78, eIF2α, XBP1, ATF6 and CHOP) were analyzed using immunofluorescence, western blot and quantitative real time PCR at different time points (8 h, 14 h and 24 h) after the various treatments. Placental ER stress reported in preeclamptic women and studies in the past few couple of years indicated that the oxidative and ER stresses are closely linked events in cell homeostasis and apoptosis^29,30^. UPR comes into role before the induction of ER stress to sustain proteostasis of cellular ambience. The master regulator of UPR (GRP78) normally (in the absence of stress) binds to N-terminus of ER stress sensors (PERK, IRE1 and ATF6) of ER^31,32,33^. However in case, when threshold of stimulus can’t be coped up, the GRP78 gets dissociated and binds preferentially with misfolded proteins^34^. Dissociation of GRP78 activates all the three arms (PERK, IRE1 and ATF6) of ER stress pathway following simultaneous autophosphorylation and oligomerization of IRE1 and PERK, and mobilization of ATF6 to the Golgi for activation^35^. In the present study expression of GRP78 protein was detected as early as 8 h, reached the peak at 14 h and it started diminishing after 14 h when BeWo cells were exposed to sera having low sFlt-1 (normotensive sera) concentration (Figure: 2a,2b). Its expression was however, further upregulated when BeWo cells were administered with normotensive sera along with re-sFlt-1 as compared to NT sera alone at all the time points (8 h, 14 h, 24 h) (Figure 7a). Protein expression of GRP78 was further validated by qRT-PCR and we found similar trend of GRP78 expression at transcription level as well (Figure 2c,3c,7f). Thus the pattern of GRP78 expression at both protein and mRNA levels suggested an important role which may be played by sFlt-1 in the induction of ER stress. Emerging evidences indicate that the amplitude and kinetics of UPR signaling are tightly regulated at different levels, which has a direct impact on cell fate decisions^36^. Analysis of the signaling kinetics of all the three ER stress sensors (PERK, IRE1and ATF6) has substantiated that the activation of each branch can vary depending on the nature of the stimulus used to perturb ER function^36,37^. Activation of PERK branch of ER stress pathway results in the phosphorylation of eukaryotic translation initiation factor 2 subunit α (eIF2 α), thereby blocking protein translation and reducing the protein overload within the ER^38^. Adaptive response goes as long as the level of stress is below threshold, however in case of prolonged stress, eIF2α-ATF4-CHOP pathway gets activated, and subsequently cells move toward apoptotic pathway ^39^. In the PERK arm of ER stress pathway, we reported maximum expression of eIF2α protein at 14 h when BeWo cells were exposed to low sFlt-1 concentration (normotensive sera) (Figure 2a,2b). Addition of re-sFlt-1 to NT sera further enhanced the expression of eIF2α and the difference between NT sera and NT + re-sFlt-1 treated cells was statistically significant at all the time points (Figure 3a,3b,7b). mRNA expression of eIF2 α coincided with protein expression (Figure 2c,3c,7g). ER stress activates all UPR signaling pathways, thereby simultaneously producing antagonistic outputs^40^. Thus in order to address this paradox, we examined the molecular behavior of all cell fate regulators of the UPR. The opposing effects of PERK and IRE1 determine whether ER stressed cells will live or die^41^. Our study also demonstrated that although all three branches were activated upon induction of ER stress yet the behavior of individual signaling pathways varied markedly with time after the onset of stress. Upon ER stress, IRE1 RNase is activated through conformational change, autophosphorylation, and higher order oligomerization^42,43,44^. IRE1 initiates diverse downstream signaling of the UPR either through unconventional splicing of the transcription factor XBP-1 or through post transcriptional modifications via Regulated IRE1-Dependent Decay (RIDD) of multiple substrates^45,46,47,48^. In the present study, expression of XBP1 was maximum at 24 h in BeWo cells exposed to NT sera (Figure 2a,2b). Expression of XBP1 in our study matched with the study by Lin *et al*., 2007, (Lin *et al*., 2007)^41^ in terms of the adaptive or apoptotic path chosen when BeWo cells were exposed to increased amount of sFlt-1 as in case of NT sera + re-sFlt-1 treatment; which further enhanced the expression of XBP1 (Figure 5a,5b). The difference between NT sera and NT sera + re-sFlt-1 treated BeWo cells was found statistically significant at all the time points (Figure 7c). Protein expression of XBP1 was further validated at mRNA levels and results were found to be consistent (Figure 2c,3c,7h). ATF6, another regulator of ER stress signaling is the type II ER transmembrane transcription factor, has two isoforms, ATF6α and ATF6β ^49^. Out of these two, ATF6α has been extensively studied in the context of ER Stress. ATF6α transits to the Golgi on the progression of ER stress, where it is cleaved by site 1 (s1) and site 2 (s2) proteases, generating an activated b-ZIP factor^50^. The processed form of ATF6α translocates to the nucleus to activate UPR genes involved in protein folding, processing, and degradation^21,22^. In the present study, we reported maximum expression of ATF6 at 24 h when the BeWo cells were exposed to NT sera (Figure 2a,2b). Addition of re-sFlt-1 to NT sera further enhanced the expression of ATF6 and the difference between NT sera and NT sera + re-sFlt-1 treated BeWo cells was statistically significant (Figure 3a,3b,7d). The mRNA levels were found consistent with protein expression (Figure 2c,3c,7i). If the various UPR induced mechanisms fail to alleviate ER stress, both the intrinsic and extrinsic pathways for apoptosis come into role. The major player involved in the cell death response include PERK/eIF2α-dependent induction of the pro-apoptotic transcriptional factor CHOP. CHOP (CCAAT-enhancer-binding protein homologous protein) has been shown to be involved in ER stress-induced apoptosis both *in vitro* and *in vivo*. CHOP was shown to downregulate transcription of anti-apoptotic BCL-2^51^. Another study has shown that it can transcriptionally upregulate expression of pro-apoptotic BH3-only protein Bim^52^ which is a mediator of ER stress-induced apoptosis in several cell types^52,53^. In the present study, CHOP expression was maximum at 24 h in BeWo cells exposed to NT sera (Figure 2a,2b). Addition of re-sFlt-1 to NT sera further enhanced the expression of CHOP (Figure 3a,3b). The difference between only NT sera treated cells and NT sera + re-sFlt-1 treated cells was statistically significant at all the time points (Figure 7e). mRNA levels were found consistent with the protein expression (Figure 2c,3c,7j). In the present study, BeWo cells, when treated with two independent concentrations of recombinant sFlt-1 (re-sFlt-1), no significant expression of ER stress markers were seen either at protein or mRNA levels at 8 h. At 14 h, upregulation of all the ER stress markers except ATF6 and CHOP were noticed with 12ng/ml. On the other hand, expression of CHOP protein was maximum at 24 h indicating increased apoptosis with progression of time. Our results are in concurrence with that of Miyake *et al*., 2016 who examined the direct effect of sFlt-1 on two ovarian cancer cell lines^54^. This group studied the mechanism of cell injury induced by two different types of sFlt-1 administrations i.e. by exogenous administration of re-sFlt-1 to culture media in four cell lines and by transfection of LV-sFlt-1 into cells. They observed that both approaches effectively damaged the tumor cells and also had anti-angiogenic effects. The reduction in tumor volume in mice model was correlated positively to the dose of re-sFlt-1.

## Summary and conclusion

In summary, we explored an interaction between anti-angiogenic factor sFlt-1 and sensors of ER stress pathways including central regulator of unfolded protein response and shed light on the molecular mechanism underlying this interaction with the help of BeWo cells mimicking trophoblast cells. We extrapolated that when BeWo cells were treated with normotensive sera (NT sera) whose sFlt-1 concentration was increased by addition of recombinant sFlt-1 exogenously, the expression of ER stress markers (at both protein and mRNA levels) increased as compared to normotensive sera alone treated cells. Thus it could be deciphered that sFlt-1 may be one of the anti-angiogenic factors present in maternal sera which not only contributes to oxidative stress but also may cause endoplasmic reticulum stress. The GRP78 expression was observed in the beginning (usually at 8 h time point) and started diminishing with time progression in all the groups. However, CHOP expression appeared usually late (maximum at 24 h) as seen in cells of group2 (NT+re-sFlt-1), group3 (re-sFlt-1) and group4 (tunicamycin treatment). However signals of CHOP protein were either feeble or undetectable in cells of group1 (NT sera), and group 5 (untreated). The protein and mRNA levels of eIF2α, XBP1 and ATF6 could be visualized between 8 h and 24 h, i.e. the time point between GRP78 induction and appearance of CHOP signals.

## Authors Contributions

SM and RD conceived and designed the study. SM did all the experiments. PA assisted in experiment. NB provided the clinical inputs and critical suggestions. SDD assisted in statistical analysis. SM and RD wrote first draft. MKD gave a hand in re-editing the content. KL, AS, RK, NR, BV, SKG, and SS further reviewed the content. RD language edited the final draft.

## Acknowledgement

We are thankful to Dr. Ashutosh Kumar, Department of Anatomy, All India Institute of Medical Sciences, New Delhi, India for his scientific inputs for the study and review assistance for the final draft of the manuscript.

## Conflict of Interest

None

## Funding

Institute Research Grant for Intramural project; Research section, All India Institute of Medical Sciences, New Delhi-110029, India

**Table 1:**
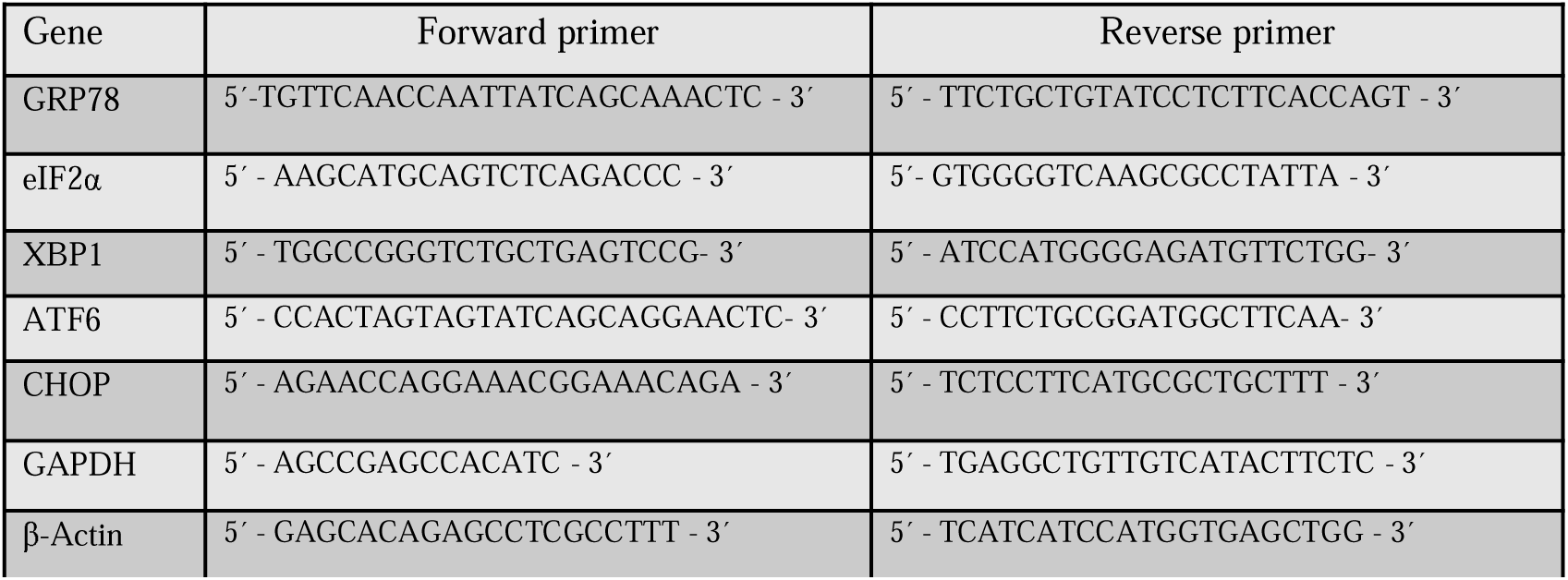
Primers: Designed by NCBI

## References

1. Hofmeyr, R., Matjila, M., Dyer, R. Preeclampsia in 2017: obstetric and anaesthesia management. Best Practice & Research Clinical Anaesthesiology., 31(1), pp. 125–138 (2017)

2. Di Renzo, G.C. The great obstetrical syndromes. J Matern Fetal Neonatal Med., 22, pp. 633–635 (2009)

3. Saftlas, A.F., Olson, D.R., Franks, A.L., Atrash, H.K., Pokras, R. Epidemiology of preeclampsia and eclampsia in the United States. Am J Obstet Gynecol., 163, pp. 460—465 (1990)

4. Khan, K.S., Wojdyla, D., Say, L., Gülmezoglu, A.M., Van Look, P.F. WHO analysis of causes of maternal death: a systematic review. Lancet., 367(9516), pp. 1066–74 (2006)

5. Maynard, S.E., Karumanchi, S.A. Angiogenic factors and preeclampsia Semin Nephrol., 31(1), pp. 33–46 (2011)

6. Siddiqui, A.H., Irani, R.A., Zhang, W., Wang, W., Blackwell, S.C., Kellems, R.E., Xia, Y. Angiotensin receptor agonistic autoantibody-mediated soluble fms-like tyrosine kinase-1 induction contributes to impaired adrenal vasculature and decreased aldosterone production in preeclampsia. Hypertension., 61(2), pp. 472–9 (2013)

7. Report of the National High Blood Pressure Education Program Working Group on High Blood Pressure in Pregnancy. Am J Obstet Gynecol., 183, pp. S1–S22 (2000)

8. MacKay, A.P., Berg, C.J., Atrash, H.K. Pregnancy-related mortality from preeclampsia and eclampsia. Obstet Gynecol., 97, pp. 533–538 (2001)

9. Charnock-Jones, D.S. Placental hypoxia, endoplasmic reticulum stress and maternal endothelial sensitisation by sFLT1 in pre-eclampsia. J Reprod Immunol., 114, pp. 81–85 (2016)

10. Yung, H.W., Korolchuk, S., Tolkovsky, A.M. Endoplasmic reticulum stress exacerbates ischemia–reperfusion-induced apoptosis through attenuation of Akt protein synthesis in human choriocarcinoma cells. FASEB J, 21, pp. 872–884 (2007)

11. Yung, H.W., Calabrese, S., Hynx, D., Hemmings, B.A., Cetin, I., Charnock-Jones, D.S., Burton, G.J. Evidence of placental translation inhibition and endoplasmic reticulum stress in the etiology of human intrauterine growth restriction. Am J Pathol., 173(2), pp. 451–62 (2008)

12. Wang, M., Kaufman, R.J. Protein misfolding in the endoplasmic reticulum as a conduit to human disease. Nature., 529, pp. 326–335 (2016)

13. Cao, S.S. Endoplasmic Reticulum Stress and Unfolded Protein Response in Inflammatory Bowel Disease. Inflammatory Bowel Diseases., 21(3), pp 636–644 (2015)

14. Hetz, C., Chevet, E., Oakes, S.A. Proteostasis control by the unfolded protein response. Nature Cell Biology., 17, pp. 829–838 (2015)

15. Jones, C.J., Fox, H. An ultrastructural and ultrahistochemical study of the human placenta in maternal pre-eclampsia. Placenta., 1(1), pp. 61–76 (1980)

16. Redman, C.W., Sargent, I.L. Latest advances in understanding preeclampsia. Science., 308, pp. 1592e4 (2005)

17. Redman, C.W. The endoplasmic reticulum stress of placental impoverishment. Am J Pathol., 173, pp. 311e4 (2008)

18. Burton, G.J., Yung, H.W., Cindrova-Davies, T., Charnock-Jones, D.S. Placental endoplasmic reticulum stress and oxidative stress in the pathophysiology of unexplained intrauterine growth restriction and early onset preeclampsia. Placenta., 30, pp. S43e8 (2008)

19. Lian, I.A., Loset, M., Mundal, S.B., Fenstad, M.H., Johnson, M.O., Eide, I.P. Increased endoplasmic reticulum stress in decidual tissue from pregnancies complicated by fetal growth restriction with and without pre-eclampsia. Placenta., 32, pp. 823e9 (2011)

20. Thilaganathan, B. Preeclampsia is not a placental disorder. Pregnancy Hypertension., 9, pp. 2 (2017)

21. Burton, G.J., Yung, H.W., Cindrova-Davies, T. Placental endoplasmic reticulum stress and oxidative stress in the pathophysiology of unexplained intrauterine growth restriction and early onset preeclampsia. Placenta., 30, pp. S43–S48 (2009)

22. Cerdeira, A.S. Karumanchi, S.A. Angiogenic Factors in Preeclampsia and Related Disorders. Cold Spring Harb Perspect Med., 2(11), pp. a006585. (2012)

23. Olsson, A.K., Dimberg, A., Kreuger J, Claesson-Welsh L. VEGF receptor signalling - in control of vascular function. Nat Rev Mol Cell Biol., 7(5), pp. 359–71. (2006)

24. Kweider, N., Huppertz, B., Wruck, C.J., Beckmann, R., Rath, W., Pufe, T., and Kadyrov, M. A. Role for Nrf2 in Redox Signalling of the Invasive Extravillous Trophoblast in Severe Early Onset IUGR Associated with Preeclampsia. PLOS., 7(10), pp. e47055 (2012)

25. Yung, H.W., Calabrese, S., Hynx, D., Hemmings, B.A., Cetin, I., Charnock-Jones, D.S., and Burton, G.J. Am J Pathol., 173(2), pp. 451–62. (2008)

26. Chaiworapongsa, T., Romero, R., Espinoza, J., Bujold, E., Mee Kim, Y., Gonçalves, L.F., Gomez, R., and Edwin, S. Urinary Placental Growth Factor in Pregnancies Complicated by Preeclampsia. Journal of Clinical and Diagnostic Research., 4626, pp. 2295. (2012)

27. Jiang, Z., Zou, Y., Ge, Z., Zuo, Q., Huang, S.Y., and Sun, L. A Role of sFlt-1 in Oxidative Stress and Apoptosis in Human and Mouse Pre-Eclamptic Trophoblasts. Biology of Reproduction., 93(3), pp. 1–7. (2015)

28. Varughese, B., Luthra K., Kumar, R., Bhatla N., Dwivedi S.N., and Dhingra, R. Urinary placental growth factor in preeclampsia. Journal of Clinical and Diagnostic Research., 6(6), pp. 929–932. (2012)

29. Burton, G.J., Yung, H.W., Cindrova-Davies, Charnock-Jones, T. Placental Endoplasmic Reticulum Stress and Oxidative Stress in the Pathophysiology of Unexplained Intrauterine Growth Restriction and Early Onset Preeclampsia. Placenta., 30, pp. 43—48 (2009)

30. Cao, S.S., Kaufman, R.J. Endoplasmic Reticulum Stress and Oxidative Stress in Cell Fate Decision and Human Disease. Antioxid Redox Signal., 21(3), pp. 396–413 (2014)

31. Bertolotti, A., Zhang, Y., Hendershot, L.M., Harding, H. P., and Ron, D. Dynamic interaction of BiP and ER stress transducers in the unfolded-protein response. Nature Cell Biol., 2, pp. 326–332 (2000)

32. Shen, J., Chen, X., Hendershot, L., Prywes, R. ER stress regulation of ATF6 localization by dissociation of BiP/GRP78 binding and unmasking of Golgi localization signals. Dev. Cell., 3, pp. 99–111 (2002)

33. Kimata, Y. Genetic evidence for a role of BiP/Kar2 that regulates Ire1 in response to accumulation of unfolded proteins. Mol. Biol. Cell., 14, pp. 2559–2569 (2003)

34. Sano, R., Reed, J.C. ER stress-induced cell death mechanisms. Biochim Biophys Acta., 1833(12), pp. 3460–3470 (2008)

35. Todd, D.J., Lee, A.H., Glimcher, L.H. The endoplasmic reticulum stress response in immunity and autoimmunity. Nat Rev Immunol., 8(9), pp. 663–74 (2008)

36. DuRose, J.B., Tam, A.B., Niwa, M. Intrinsic Capacities of Molecular Sensors of the Unfolded Protein Response to Sense Alternate Forms of Endoplasmic Reticulum Stress. Mol Biol Cell., 17(7), pp. 3095–3107 (2006)

37. Yoshida, H. A time-dependent phase shift in the mammalian unfolded protein response. Dev. Cell., 4, pp. 265–271 (2003)

38. Szegezdi, E., Logue, S. E., Gorman, A. M., Samali, A. Mediators of endoplasmic reticulum stress-induced apoptosis. EMBO reports., 7, pp. 880–885 (2006)

39. Han, J., Back, S.H., Hur, J., Lin, Y.H., and Kaufman, R.J. ER-stress-induced transcriptional regulation increases protein synthesis leading to cell death. Nat Cell Biol., 15(5), pp. 481–490 (2013)

40. Lin, J.H., Li, H., Yasumura, D., Cohen, H.R., and Walter, P. IRE1 signaling affects cell fate during the unfolded protein response. Science., 318(5852), pp. 944–9 (2007)

41. Kanekura, K., Ma, X., Murphy, J.T., Zhu, L.J., Diwan, A., and Urano, F. IRE1 prevents endoplasmic reticulum membrane permeabilization and cell death under pathological conditions. Sci Signal., 8(382), pp. ra62 (2015)

42. Prischi, F., Nowak, P.R., Carrara, M., Ali, M.M.U. Phosphoregulation of Ire1 RNase splicing activity. Nature Communications., 5, pp. 3554 (2014)

43. Aragon, T. Messenger RNA targeting to endoplasmic reticulum stress signalling sites. Nature., 457, pp. 736–740 (2009)

44. Korennykh, A. V. The unfolded protein response signals through high-order assembly of Ire1. Nature., 457, pp. 687–693 (2009)

45. Hetz, C., Martinon, F., Rodriguez, D., Glimcher, L. H. The unfolded protein response: integrating stress signals through the stress sensor IRE1α. Physiol. Rev., 91, pp. 1219–1243 (2011)

46. Hollien, J. Regulated Ire1-dependent decay of messenger RNAs in mammalian cells. J. Cell Biol., 186, pp. 323–331 (2009)

47. So, J.S. Silencing of Lipid Metabolism Genes through IRE1 alpha-Mediated mRNA Decay Lowers Plasma Lipids in Mice. Cell Metabolism., 16, pp. 487–499 (2012)

48. Sakaki, K. RNA surveillance is required for endoplasmic reticulum homeostasis. Proc Natl Acad Sci US A., 109, pp. 8079–8084 (2012)

49. Yung, H.W., Korolchuk, S., Tolkovsky, A.M. Endoplasmic reticulum stress exacerbates ischemia–reperfusion-induced apoptosis through attenuation of Akt protein synthesis in human choriocarcinoma cells. FASEB J. 21, pp. 872–884 (2007)

50. Yung, H.W., Calabrese, S., Hynx, D., Hemmings, B.A., Cetin, I., Charnock-Jones, D.S., and Burton, G.J. Am J Pathol., 173(2), pp. 451–62 (2008)

51. Walter, P., Ron, D. The unfolded protein response: from stress pathway to homeostatic regulation. Science., 334, pp. 1081–1086 (2011)

52. McCullough, K.D., Martindale, J.L., Klotz, L.O., Aw, T.Y., and Holbrook, N.J. Gadd153 sensitizes cells to endoplasmic reticulum stress by down-regulating Bcl2 and perturbing the cellular redox state. Mol. Cell. Biol., 21, pp. 1249–1259 (2001)

53. Puthalakath, H., O'Reilly, L.A., Gunn, P., Lee, L., Kelly, P.N., Huntington, N.D., Hughes, P.D., Michalak, E.M., McKimm-Breschkin, J., Motoyama, N., Gotoh, T., Akira, S., Bouillet, P., Strasser, A. ER stress triggers apoptosis by activating BH3-only protein Bim. Cell., 129, pp. 1337–1349 (2007)

54. Miyake, T., Kumasawa, K., Sato, N., Takiuchi, T., Nakamura, H., and Kimura T. Soluble VEGF receptor 1 (sFLT1) induces non-apoptotic death in ovarian and colorectal cancer cells. Scientific Reports., 6, pp. 24853 (2016)

